# Measuring the tolerance of the genetic code to altered codon size

**DOI:** 10.1101/2021.04.26.441066

**Authors:** E. DeBenedictis, D. Söll, K. Esvelt

## Abstract

Protein translation using four-base codons occurs in both natural and synthetic systems. What constraints contributed to the universal adoption of a triplet-codon, rather than quadruplet-codon, genetic code? Here, we investigate the tolerance of the *E. coli* genetic code to tRNA mutations that increase codon size. We found that tRNAs from all twenty canonical isoacceptor classes can be converted to functional quadruplet tRNAs (qtRNAs), many of which selectively incorporate a single amino acid in response to a specified four-base codon. However, efficient quadruplet codon translation often requires multiple tRNA mutations, potentially constraining evolution. Moreover, while tRNAs were largely amenable to quadruplet conversion, only nine of the twenty aminoacyl tRNA synthetases tolerate quadruplet anticodons. These constitute a functional and mutually orthogonal set, but one that sharply limits the chemical alphabet available to a nascent all-quadruplet code. Our results illuminate factors that led to selection and maintenance of triplet codons in primordial Earth and provide a blueprint for synthetic biologists to deliberately engineer an all-quadruplet expanded genetic code.

## INTRODUCTION

The genetic code is determined by a combination of tRNAs and aminoacyl tRNA synthetases (AARSs). Codons are dictated by the three bases in the center of the anticodon loop of each tRNA, which undergo Watson-Crick base pairing to an mRNA transcript during translation, enabling accurate codon recognition. The correspondence between codons and amino acids — one tRNA isoacceptor class for each canonical amino acid — is dictated by the twenty AARSs which specifically recognize bases (identity elements) in the tRNAs, and attach the cognate amino acid onto only the CCA 3’ terminus of the cognate tRNA. The aminoacylation process is exquisitely accurate, enabling high-fidelity protein synthesis^1^. Anticodon mutations frequently alter or abolish selective charging with the cognate amino acid because most AARSs rely on bases in the anticodon to identify the cognate tRNA^2^. However, certain natural anticodon mutations generate ‘suppressor’ tRNAs that insert their cognate amino acid in response to 5’-UAG-3’ stop codons using a cognate 5’-CUA-3’ anticodon^3^. Frameshift suppression, in which a quadruplet tRNA reads through a frameshift mutation, are also known to arise naturally^4,5^; i.e. qtRNA-Gly-GGGG can decode the four-base codon 5’-GGGG-3’ in mRNA transcripts using the 5’-CCCC-3’ anticodon. In the presence of efficiently aminoacylated qtRNAs, the ribosome is capable of protein translation with individual non-canonical stop or quadruplet codons within an otherwise all-triplet transcript^6^. If individual quadruplet codon translation is known to arise through simple point insertions, what functional constraints, if any, prevent the natural or synthetic evolution of an all-quadruplet genetic code?

These origin-of-life questions have newfound importance to engineering with the advent of genetic code expansion technology. An expanded all-quadruplet genetic code would offer 256 total codons, including hundreds of free codons that could be assigned to noncanonical amino acids (ncAAs), valuable chemical additions to the genetic code that enable improved protein therapeutics^7^. Most work to date has focused on incorporating ncAAs using quadruplet codons within otherwise all-triplet mRNA transcripts^8^. An all-quadruplet genetic code would presumably require the ability to incorporate the 20 canonical amino acids with quadruplet codons. Might the necessary translation components be easily derived from existing tRNAs and AARSs?

To investigate these questions, we assessed the tolerance of the genetic code to mutations that alter codon size in *E. coli*. We systematically explored whether quadruplet codon translation can arise through simple point insertions in each of the tRNA anticodon loops, or through mutation of many bases in the anticodon, then used directed evolution to determine how often additional mutations throughout the tRNA can improve translation of the resulting quadruplet qtRNAs. Finally, we characterized the fidelity quadruplet codon translation. Remarkably, we found that 12/20 isoacceptor classes of tRNAs can be readily converted to selectively charged qtRNAs, that activity is low but can often be improved by accumulating additional mutations along the sides of the anticodon loop, and that most of the resulting qtRNAs selectively incorporate a single amino acid in response to a quadruplet codon. Our results suggest that quadruplet codon translation was undoubtedly explored by primordial organisms, identifies the barriers limiting its complete adoption by natural evolution, and presents 9/20 qtRNAs necessary to create an all-quadruplet expanded genetic code.

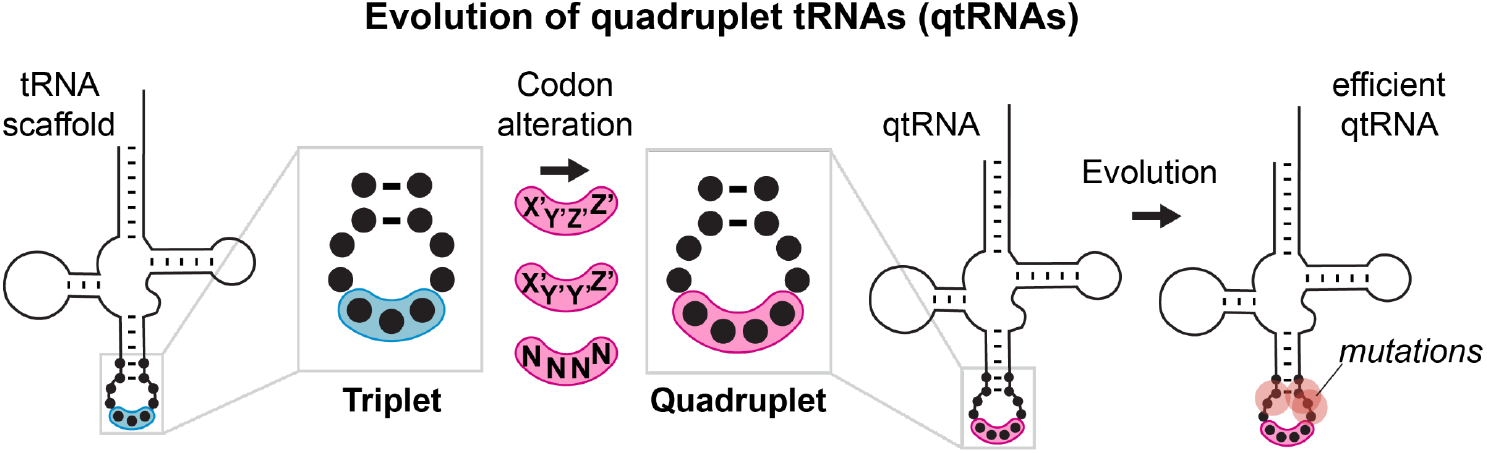
Visual abstract. We studied whether tRNAs can arise through simple changes to the anticodon followed by additional mutations accumulated during evolution.

## RESULTS

### Evolution of qtRNAs through point insertions

Many known examples of frameshift suppressors contain single point insertions in the anticodon that convert a triplet tRNA into a quadruplet tRNA (qtRNA). We initially tested whether tRNAs can evolve into qtRNAs through simple point insertions. We selected 21 endogenous *E. coli* tRNAs, one cognate tRNA for each canonical amino acid and initiator methionine (Table S1), to serve as ‘scaffolds’ into which we introduce anticodon point insertions. We tested two point insertion locations: if the original tRNA decodes the triplet codon “XYZ”, we created a qtRNA that decodes “XYZZ” and a qtRNA that decodes “XYYZ” (Figure 1A). These anticodon patterns preserve the nature of the bases in the anticodon loop, which most AARSs use to recognize the cognate tRNA^2^, and are found in known qtRNAs, such as sufD^4^ (qtRNA^Gly^_GGGG_) and sufG^9^ (qtRNA^Gln^_CAAA_). Throughout, for ease of comparison to standard triplet codon tables, we use qtRNA^three letter scaffold^_four letter DNA codon_ nomenclature to refer to qtRNAs; e.g. a serine qtRNA bearing a 5’-UCUA-3’ anticodon is referred to as qtRNA^Ser^_TAGA_.

**Figure 1:**
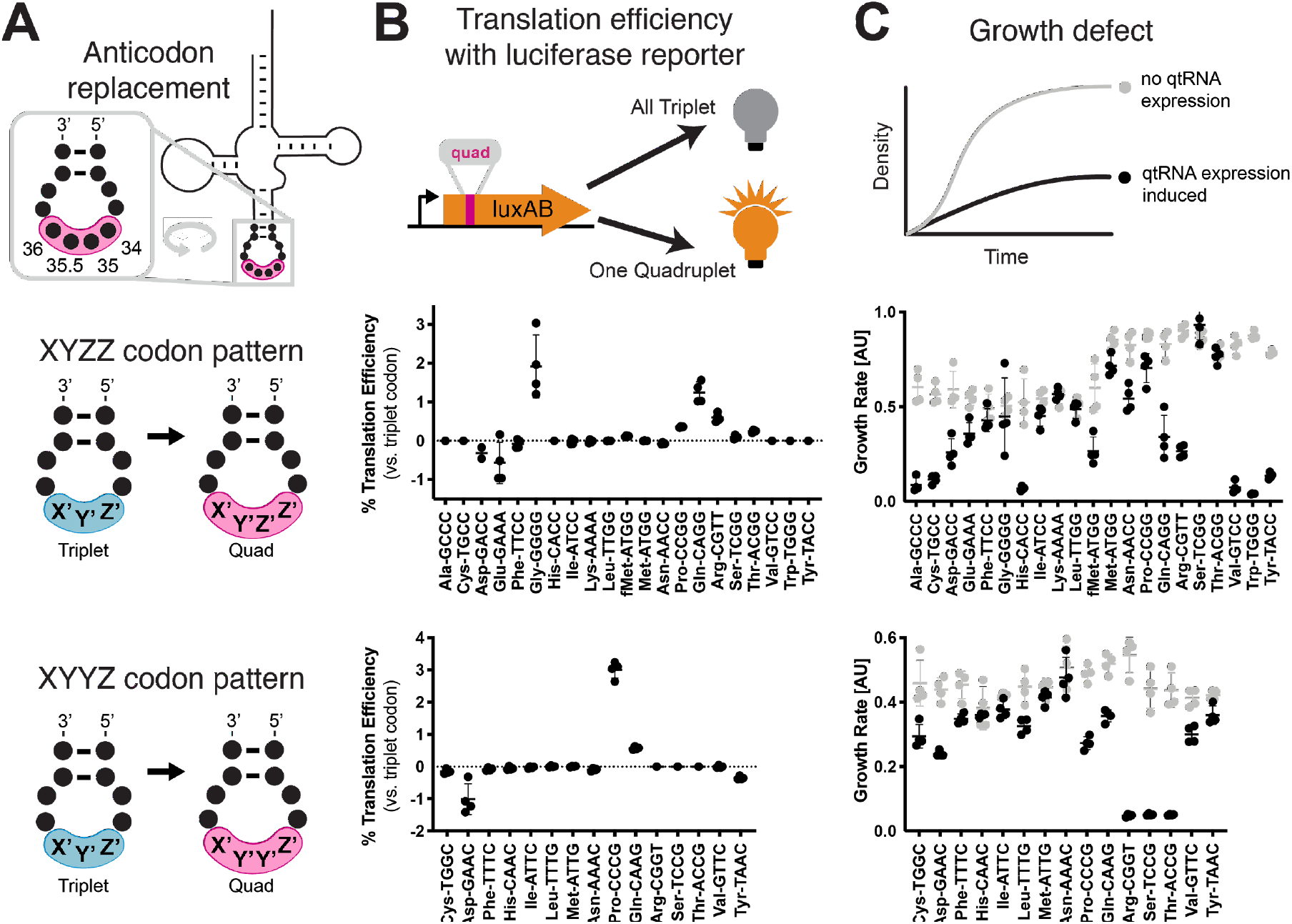
**A)** We measured qtRNAs that might arise through point insertions in the anticodon loop. Each qtRNA is based upon a tRNA from the *E coli* genome that serves as a ‘scaffold’ (Table S1). We tested two quadruplet codon patterns: a tRNA decoding the triplet codon “XYZ” to a qtRNA decoding the quadruplet codon “XYZZ” or “XYYZ”. Note that in instances in which XYZZ and XYYZ are the same, the qtRNA is depicted on the XYZZ graph. For clarity, we use “qtRNA”-”three letter scaffold”-”four letter codon” nomenclature to refer to qtRNAs; e.g. a Serine qtRNA bearing a 5’-UCUA-3’ anticodon that recognizes 5’-UAGA-3’ in mRNA transcripts is referred to as qtRNA^Ser^_TAGA_. **B)** We measured these qtRNAs using a luciferase readthrough assay. In the presence of a functional qtRNA generates full-length luxAB transcript and luminescence, while nonfunctional qtRNAs generate a truncated luxAB transcript and no luminescence. Measurements are taken kinetically and normalized to culture density. Efficiency is reported relative to luminescence produced by a WT, all triplet luciferase transcript. Note that fMet qtRNAs are measured with a luciferase reporter bearing a quadruplet codon at residue 1; all others are measured with a quadruplet codon at residue 357 of luxAB. **C)** Expression of qtRNAs can be toxic. Here we report the density difference between cultures where qtRNA expression had been induced (black) or suppressed (grey).

We used two techniques to characterize these qtRNAs. First, to measure the quadruplet codon translation efficiency, we used a luciferase readthrough assay. This reporter contains a single quadruplet codon at permissive residue 357 of *luxAB*; failure to decode the quadruplet codon leads to premature termination, whereas successful four base decoding results in full-length luxAB translation and luminescence (Figure 1B).

Seven of the twenty “XYZZ”-decoding qtRNAs and two of the “XYYZ”-decoding qtRNAs functionally decode a quadruplet codon during translation. The frequent functionality of the XYZZ codon pattern may be due to flexibility in synthetase recognition at the third position of the codon, which is frequently a wobble base pair. Next, we quantified the toxicity of qtRNA expression by comparing the growth of cultures with qtRNA expression induced or suppressed (Figure 1C). The qtRNAs fall into several categories: those that exhibit no fitness defect such as the naturally occurring and highly functional qtRNA^Gly^_GGGG_; those that exhibit severe fitness defects and effectively halt bacterial growth upon induction such as qtRNA^Ala^_GCCC_, and those that moderately slow bacterial growth. The translation efficiency and growth defect of qtRNAs depends upon more than just the interaction with the cognate AARS: AlaRS, LeuRS, and SerRS are all synthetases that do not interact with the anticodon loop of their cognate tRNA, but qtRNAs derived from these scaffolds exhibit a range of behaviors depending upon the new anticodon. For example, qtRNA^Ser^_TCCG_ exhibits high growth defect, while qtRNA^Ser^_TCGG_ exhibits no growth defect and modest quadruplet translation efficiency. Together, this data shows that a third of the twenty isoacceptor classes have access to point insertions that create modestly functional quadruplet codon translation, however, other point insertions create qtRNAs that do not functionally decode quadruplet codons or incur large growth defects when expressed.

### Evolution of qtRNAs through anticodon replacement

Next, we tested whether tRNAs could evolve into qtRNAs through whole anticodon replacement, as might occur during recombination or more intense mutagenesis. Antibiotic selection markers have previously been used to identify functional qtRNAs^10^. We applied an equivalent approach based on the use of M13 bacteriophage tail fiber pIII as a selection marker^11^. For each of the 20 representative *E. coli* tRNA scaffolds used above, we created a 256-member qtRNA library containing degenerate anticodons (Figure 2A) and selected for functional qtRNAs from these libraries (Figure 2B) to identify qtRNAs that decode any of eight quadruplet codons of interest (Figure 2C). High final phage titers after selection indicate the presence of a functional qtRNA, so we used phage titer to guide us in selecting 69 putative qtRNAs for further characterization (Figure 2D). Using Sanger sequencing, we determined the anticodon of each highly selected variant. We found that for most variants, the qtRNA agrees with the quadruplet codon in the reporter at all four positions, or the first three positions of the codon (Figure 2E), in agreement with previous findings on quadruplet codon crosstalk with fourth-base mismatches^12^. We found that qtRNAs identified by this phage-based assay are also functional in the luciferase assay. These results demonstrate that most tRNA scaffolds are capable of supporting quadruplet codon translation through whole-anticodon replacement.

**Figure 2:**
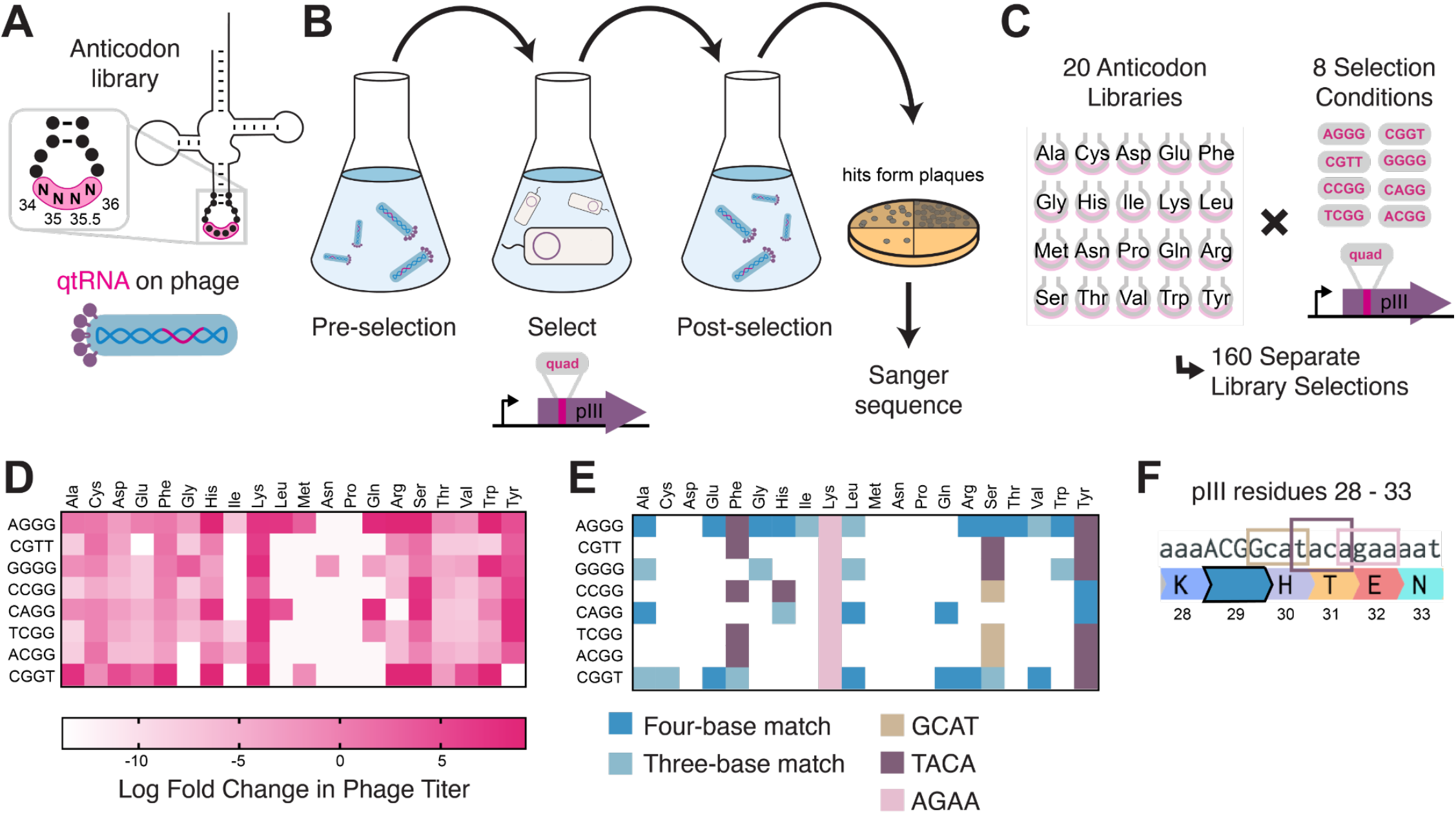
**A)** We created qtRNA libraries using degenerate primers to randomize the four bases in the anticodon. The qtRNA library is expressed from a ΔpIII M13 bacteriophage. **B)** We selected these libraries by challenging the phage to infect and propagate in bacteria that require a quadruplet codon to be translated in order to produce a functional version of the essential phage gene, pIII. Plaques from selected libraries were Sanger sequenced. **C)** We crossed each library with each pIII-based reporter for a total of 160 separate library selections. **D)** The log fold change in phage titer when comparing the post-selection population to the pre-selection population. **E)** For every filled square, we sequenced two plaques in order to determine the anticodon identity. Results include instances in which the qtRNAs match the reporter at all four positions, or that match with the first three bases of the codon (shades of blue). Additionally, we show instances in which qtRNAs were discovered that suppress a different quadruplet codon near residue 29. **F)** Although we intend for the phage to suppress the quadruplet codon located at permissive residue 29 of pIII, in several instances the selection identified qtRNAs that suppress a nearby quadruplet codon instead.

In addition to the expected anticodons, we found that some qtRNAs instead matched quadruplet codons that appear nearby in the sequence context of the reporter (Figure 2F). We noticed that the unexpected codons often bear similarity to the qtRNA’s original codon; i.e. qtRNA^Lys^_AGAA_ is highly similar to the scaffold’s original AAA codon, and qtRNA^Tyr^_TACA_ and qtRNA^Phe^_TACA_ are similar to TAC and TTC respectively. The emergence of these anticodons suggests that these isoacceptor classes favor quadruplet codons that are related to their natural triplet codon, and demonstrates that qtRNA evolution depends upon the sequence context of relevant ORFs.

Together, these experiments identified functional qtRNAs involving four or fewer mutations for 18/20 isoacceptor classes. For the remaining two isoacceptor classes, Met and Asn, we systematically tested additional codons and found that qtRNA^Met^_AGGG_ and qtRNA^Asn^_AGGA_ both exhibit weak quadruplet codon translation (Figure S1). Therefore, every isoacceptor class can give rise to qtRNAs capable of decoding quadruplet codons during protein translation.

### Directed evolution of qtRNAs

Having found that qtRNAs that functionally decode quadruplet codons can arise generally through just a few mutations, we sought to understand other factors that may prevent more widespread use of quadruplet codons. Many qtRNAs were quite inefficient in translation: the presence of a single quadruplet codon in an mRNA transcript can reduce total protein yield to less than 3% relative to an all-triplet mRNA. Mutations at the anticodon loop sides of the qtRNAs have been observed to improve translation efficiency for TAGA-qtRNAs^11,13,36^. Triplet tRNAs are known to exhibit patterns that relate the bases in the ALS to the bases in the anticodon itself^14^, and similarly benefit from ALS mutations after anticodon replacement^15–17^. In some cases, mutations in this area can alter qtRNA charging; in others they improve quadruplet translation efficiency without altering the qtRNA’s interaction with the cognate AARS^36^. We hypothesized that qtRNAs in general require mutations at bases 32, 37 and 38 to better accommodate a new codon, and that this requirement may present a key barrier preventing the natural evolution of efficient qtRNAs.

To test this hypothesis experimentally, we selected 41 promising qtRNAs, including at least one qtRNA for every unique scaffold, and cloned a library containing degenerate nucleotides at bases 32, 37, and 38 (Figure 3A). We selected functional members of these libraries using the pIII-based selection and used next generation sequencing to characterize the abundance of each library member before and after selection (Figure 3B).

**Figure 3:**
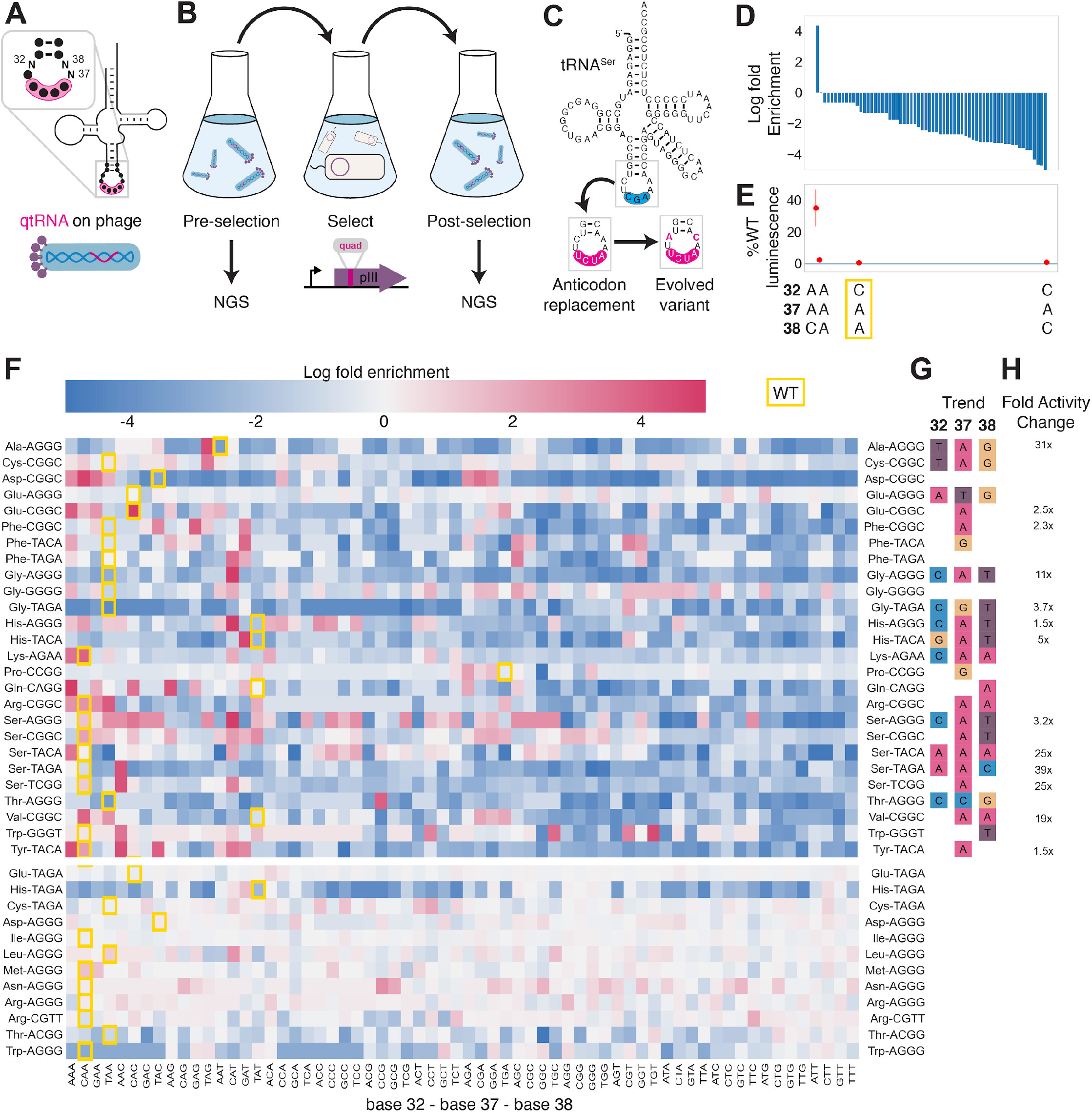
**A)** We created qtRNA libraries using degenerate primers to randomize positions 32, 37 and 38 of the anticodon. The qtRNA library is expressed from a ΔpIII M13 bacteriophage. **B)** We selected these libraries by challenging the phage to infect and propagate in bacteria that require a quadruplet codon to be translated in order to produce a functional version of the essential phage gene, pIII. Libraries were NGS sequenced to >10x library size before and after selection. **C)** tRNA^Ser^_TCG_ is known to be a scaffold for the functional qtRNA^Ser^_TAGA_ after anticodon replacement alone. Additional mutations to the sides of the anticodon loop are known to improve quadruplet codon translation efficiency. **D)** Log fold enrichment of the population abundance and **E)** translation efficiency as measured by a luciferase readthrough assay of the 64 possible combinations of nucleotides at positions 32, 37 and 38. **F)** Log fold enrichment of all 41 qtRNA libraries for each of the 64 library members. Libraries are separated by those that exhibit abundance changes during selection (above) from those that exhibit no significant abundance changes (below). **G)** For each library, the trend in nucleotide preference for each position is listed. **H)** Nucleotide preferences for select libraries were measured by cloning a qtRNA variant and measuring it using a luciferase readthrough assay. The fold improvement in activity over the wildtype values of base 32, 37 and 38 are listed. In e) and f), the original identities of bases 32, 37, and 38 found in the wildtype triplet tRNA scaffold are boxed in gold.

We began by assessing results for the well-studied qtRNA^Ser^_TAGA_, which is known to have an improved variant, qtRNA^Ser^_TAGA_-32A-38C, that exhibits improved quadruplet codon translation but unaltered, selective aminoacylation with serine^36^ (Figure 3C). Of the 64 possible combinations of DNA bases at positions 32, 37, and 38, the single library member A32 A37 C38 is enriched four log fold above all other variants (Figure 3D). We measured several qtRNA^Ser^_TAGA_ variants that correspond to different levels of enrichment using a luciferase readthrough assay, and confirmed that the strongly enriched variant exhibits more efficient quadruplet decoding than deenriched variants (Figure 3E).

Next, we applied the same procedure to quantify fold enrichment for the other 40 qtRNA libraries (Figure 3F). We identified 15 libraries that exhibit no selective pressure, 12 libraries that strongly enrich a single library member, like qtRNA^Ser^_TAGA_, and an additional 11 libraries exhibit strong enrichment for library members with a specific base at one or two positions, but not all three (Figure 3G). For several of these libraries, we used a luciferase readthrough assay to measure the fold change activity when mutations to bases 32, 37 and 38 are introduced. In most cases, introduction of these mutations substantially increases quadruplet codon translation efficiency (Figure 3H).

We were curious whether there are overall trends in the identity of optimal anticodon loop sides for efficient quadruplet codon translation. The optimal library member is not determined by the scaffold or codon independently, indicating that the mechanism by which these mutations improve quadruplet codon translation does not improve the qtRNA’s interaction with its respective AARS. Amongst the libraries there was a prominent preference for A37, a base known to be associated with reading frame maintenance^18^. Additionally, C32 A37 T38 and T32 A37 G38 appear often amongst libraries that exhibit strong preference for a single library member. The presence of modified nucleotides is especially important at two sites: position 34, the first base of the anticodon, and position 37, the nucleotide downstream of the anticodon^18,20^. The location of ‘identity elements’ for some of these modifying enzymes within the anticodon loop sides has been implied^20^. Mutation of these bases may improve the RNA modification of the anticodon loop, which are essential for many tRNA functions^19^.

### Which amino acid(s) do qtRNAs incorporate during translation?

Having established that anticodon modification can often produce tRNAs that decode four-base codons, we sought to determine which amino acid these qtRNAs incorporate during translation. To do so, we translated *sfGFP* mRNA containing a quadruplet codon at permissive residue 151 in the presence of a qtRNA. We then used mass spectral analysis to determine the nature and occupancy of amino acid 151 in the resulting protein^17^ for qtRNAs based on the 20 distinct tRNA scaffolds (Figure 4).

**Figure 4:**
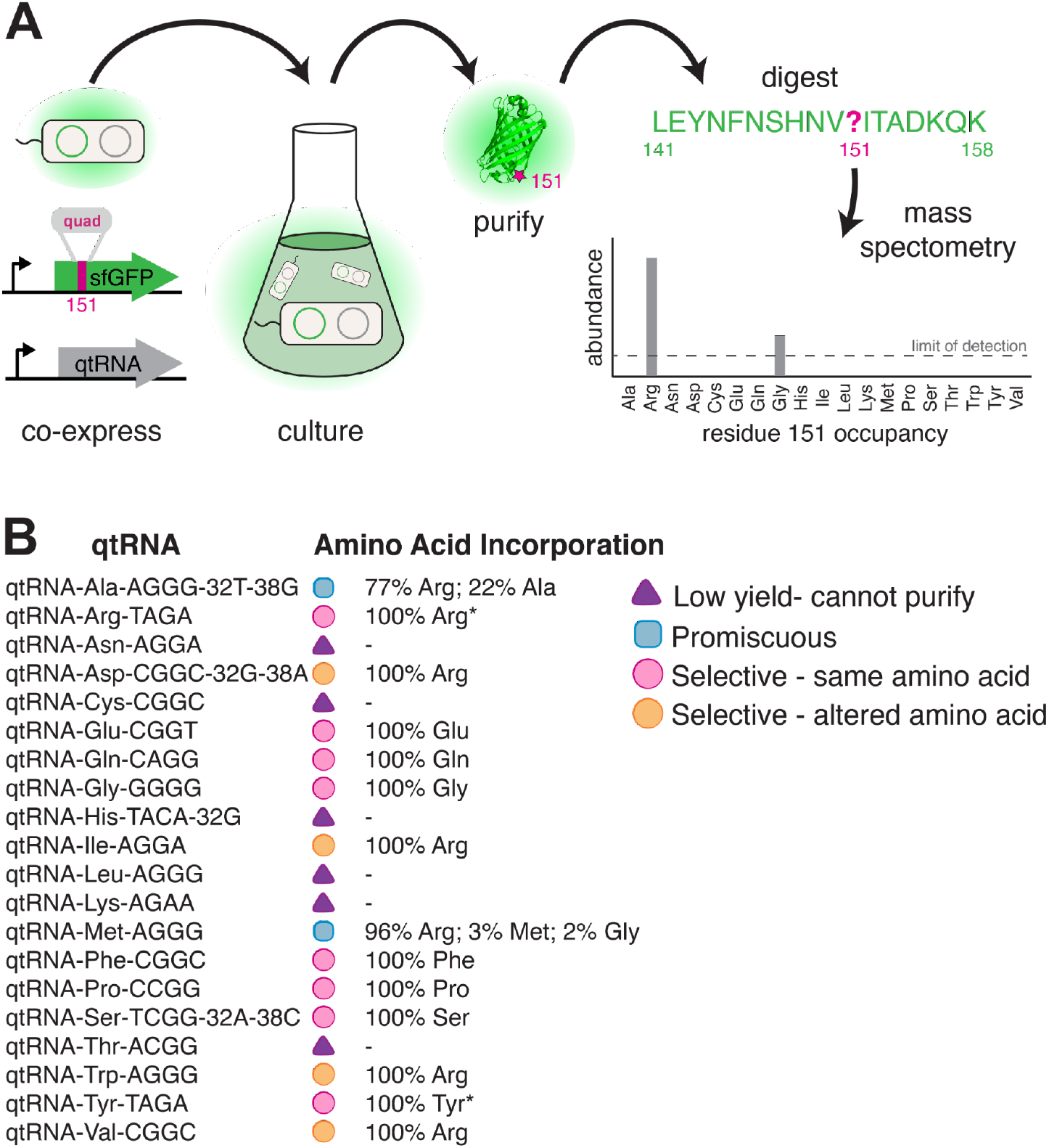
Characterization of amino acid incorporation by qtRNAs. **A)** We characterized the amino acid incorporated during translation by co-expressing a qtRNA and a sfGFP-151-quad transcript, purifying the resulting GFP, and analyzing the occupancy of residue 151 using mass spectrometry. **B)** Results of applying this pipeline to at least one qtRNA based on each of the 20 canonical scaffolds. For some qtRNAs, yield was too low to allow for charging characterization even when purified at 1L scale (Table S2). Charging of (*) qtRNA^Arg^_TAGA_ and qtRNA^Tyr^_TAGA_ were first reported in [36]

We were unable to characterize the acylation properties of six qtRNAs due to low sfGFP purification yield, even when purified at 1-L scale (Table S2) (qtRNA^Asn^, qtRNA^Cys^, qtRNA^Leu^, qtRNA^Lys^, qtRNA^Met^, and qtRNA^Thr^). These qtRNAs would be unlikely to be selected in a natural system due to their inability to produce full-length protein efficiently. Eight qtRNAs are selectively acylated with the amino acid cognate to the original scaffold (qtRNA^Arg^, qtRNA^Gln^, qtRNA^Glu^, qtRNA^Gly^, qtRNA^Phe^, qtRNA^Pro^, qtRNA^Ser^, qtRNA^Tyr^). Four qtRNAs selectively incorporate Arg, rather than the amino acid cognate to the scaffold (qtRNA^Asp^, qtRNA^Ile^, qtRNA^Trp^, qtRNA^Val^). Two qtRNAs that are charged with the cognate amino acid as well as promiscuously charged with Arg (qtRNA^Ala^ and qtRNA^Met^). All 14 of these qtRNAs might be selected by a natural system to rescue a frameshift mutation. The 12 qtRNAs that are selectively charged might be further selected as components of an all-quadruplet genetic code.

These data highlight the plasticity of AARS recognition for altered codons, both in nucleobase composition and size. The presence of a quadruplet anticodon presumably distorts the structure of the anticodon binding domain, which is a major identity element for many AARSs^2^. How much distortion will the enzyme functionally accept, and/or will recognition of the additional base lead to mischarging with a different amino acid? Of the eight qtRNAs that retained the cognate specificity, tRNA^Gly^ and tRNA^Pro^ are known as naturally occurring functional frameshift suppressors^4,21^; we confirm that they are selectively charged by the cognate AARS. Together with tRNA^Gln^, our results show that *E. coli* GlnRS, GlyRS and ProRS recognize the first three bases of the quadruplet anticodon, and are capable of charging qtRNAs despite the increased anticodon size. Correct charging of qtRNAs derived from qtRNA^Ser^ is expected, as *E. coli* SerRS does not interact with the anticodon^22^. A striking result is the correct charging of qtRNA^Glu^_CGGT_ and qtRNA^Phe^_CGGC_ by their respective *E. coli* tRNA synthetases, as their triplet anticodon sequence is unlike that of the quadruplet anticodon. However, in both cases all other known critical identity elements^2^ are present in the tRNA scaffold. This means that the major recognition feature, the anticodon, can be outweighed by the sum of the other identity elements, creating in some cases an avenue for anticodon evolution.

Finally, why did we observe acylation of multiple qtRNAs with arginine? ArgRS is responsible for synthesis of a family of Arg-tRNAs needed for the recognition of six codons, including tRNAs that differ at both the first and last position of the anticodon. As a consequence, the identity of just one anticodon position is invariant (C35). Examination of the qtRNAs charged with Arg (Table 4B) shows that all satisfy this anticodon identity element, causing rampant promiscuous charging with Arg, even for qtRNAs that lack other known argRS identity elements such as A20 such as qtRNA^Asp^_CGGC_ and qtRNA^Val^_CGGC_. For this reason, the presence of promiscuous ArgRS substantially increases the probability that tRNA point insertions in diverse scaffolds will result in an aminoacylated qtRNA.

Taken together, we found that in the majority of cases, qtRNAs exhibit properties that would render them evolutionarily favored building blocks: they are selectively charged by a single amino acid and efficiently incorporate that amino acid in response to their quadruplet codon.

### Trends in nascent qtRNA evolution

In total, we characterized 116 different qtRNAs based on 20 tRNA scaffolds that decode 20 unique quadruplet codons (Figure 5A). This vastly expands the total number of known qtRNAs, the diversity of triplet tRNA scaffolds they are based upon, and the diversity of quadruplet codons they recognize beyond what has previously been reported. We found that 60 out of 109 are functional, that is, they generate increased luminescence upon induction of qtRNA expression. In total, every tRNA scaffold we tested is capable of supporting quadruplet codon translation given appropriate four-base codon choice.

**Figure 5:**
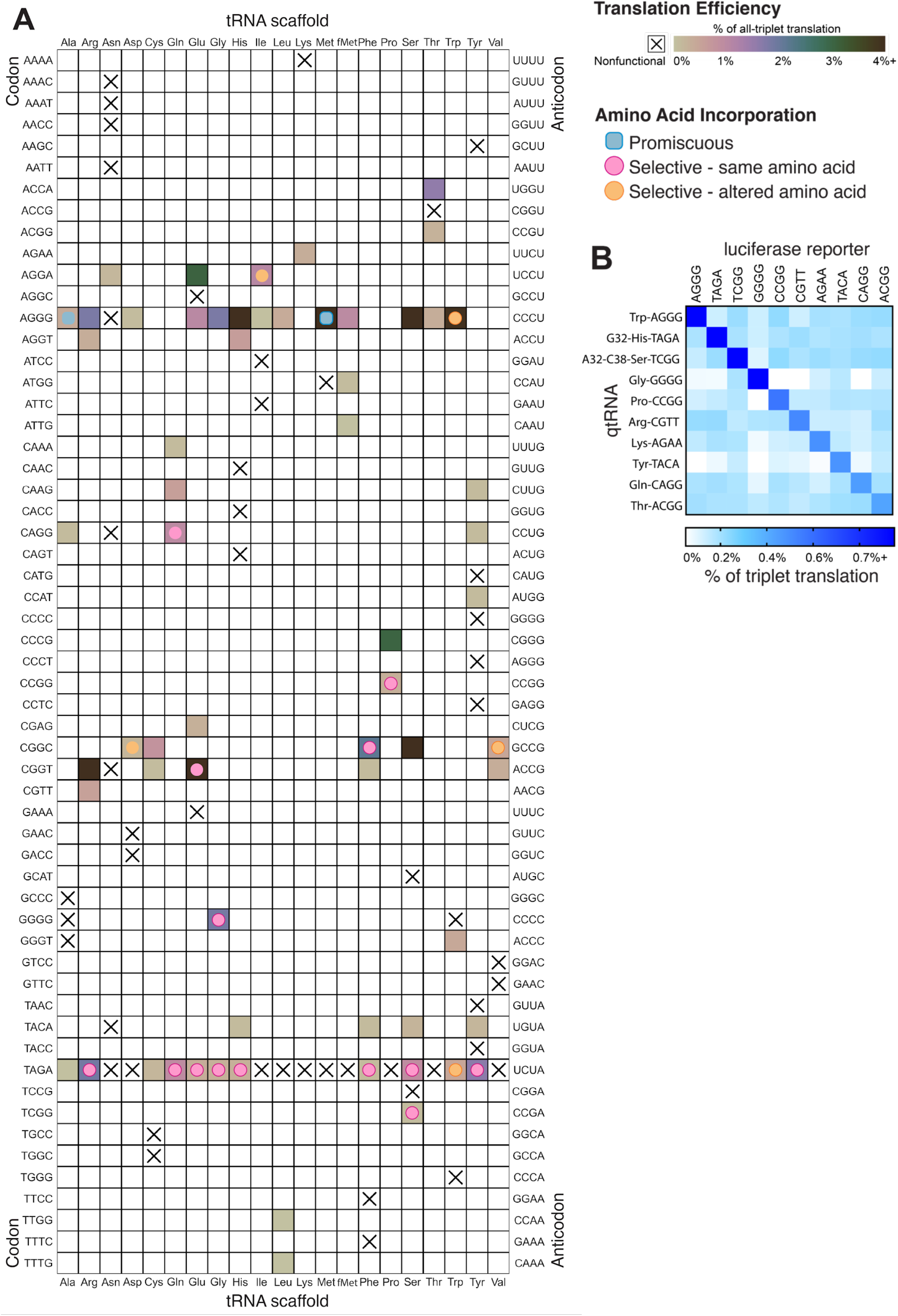
**A)** Compiled results of quantifying qtRNA translation efficiency (luciferase redthrough assay) and charging (GFP mass spectrometry). The qtRNA scaffold (columns) and codon (rows) are indicated. The translation efficiency is indicated on each qtRNA that could be measured, or marked as “nonfunctional”, meaning the qtRNA exhibits too strong a growth defect to measure, or does not produce a measurable increase luminescence upon induction of qtRNA expression. Translation efficiency is measured as a percent of all-triplet translation (see Methods). Note that fMet qtRNAs are measured with a luciferase reporter bearing a quadruplet codon at residue 1; all others are measured with a quadruplet codon at residue 357 of luxAB. Charging results are indicated on the table for qtRNAs derived from each scaffold-codon pair; measured qtRNAs may contain additional mutations; see Table S2. **B)** A miniature all-quadruplet genetic code. For each of the 10 qtRNAs (rows), we measured readthrough of a luciferase transcript containing the indicated quadruplet codon at residue 357 (columns).

Could these codon-diverse qtRNAs, if accumulated in a genome, enable all-quadruple translation? To be used as part of a genetic code, qtRNAs must be capable of operating together with minimal crosstalk. We selected 10 qtRNAs and measured crosstalk between members of this set, revealing a high degree of orthogonality: each qtRNA is specific for its own four-base codon (Figure 5B). Additionally, we found eight novel qtRNAs that selectively incorporate the same amino acid as the scaffold, despite the altered codon. Together, nine different AARSs (ArgRS, GluRS, GlnRS, GlyRS, HisRS, PheRS, ProRS, SerRS, TyrRS) faithfully and efficiently charge a cognate qtRNA, creating the opportunity for amino acid and codon-diverse all quadruplet peptide translation.

## DISCUSSION

Primordial Earth certainly sampled qtRNAs during early evolution: we found that every tRNA in *E. coli* is no more than 4 mutations away from supporting quadruplet codon translation. Indeed, single point insertions in the anticodon generate functional qtRNAs for 7/20 isoacceptor classes. The most efficient of these, such as qtRNA^Gly^_GGGG_, have been discovered as naturally occurring frameshift suppressors^4,5^, indicating that existing approaches have been successful in identifying only the most efficient qtRNAs. There are an additional 5 isoacceptor classes that can support reasonably efficient quadruplet codon translation with whole-anticodon replacement. We found that 12/20 isoacceptor classes result in qtRNAs that are selectively charged by a single amino acid, making them putative components of novel, high-fidelity genetic codes. Additionally, the diversity of codons that can be recognized by qtRNAs can easily enable codon-selective translation, as we demonstrate with a miniature all-quadruplet genetic code composed of 10 orthogonal qtRNAs based on unique isoacceptor classes. Together, these results show that qtRNAs certainly arose many times during evolution in primordial Earth, and that many of the qtRNAs were capable of amino-acid selective, efficient translation of quadruplet codons.

However, our study identifies several factors that may have prevented universal adoption of an all-quadruplet code. First, we find that some qtRNAs exhibit strong fitness defects which prevent them from being maintained and accumulated in genomes. Second, additional mutations outside the anticodon are often required for efficient translation, reducing the probability that efficient qtRNAs would arise by chance. Such mutations may affect identity elements of the tRNA modifying enzymes that chemically alter the bases surrounding the anticodon^23^; similar patterns have been observed for canonical triplet tRNAs^14^. Finally, only some AARSs are amenable to altered codon sizes for some amino acids: 12/20 modern *E. coli* AARSs are intolerant, inefficient, or promiscuous with qtRNAs. In particular, AARSs that arose later during evolution may be more finely tuned to precise anticodon base recognition and thus less tolerant of anticodon mutations or expansion. A nascent all-quadruplet code would initially have access to only the limited chemical vocabulary of the nine AARSs that are most amenable to quadruplet codon translation, decreasing the likelihood that valuable quadruplet-ORFs might arise and be selected. Finally, studies have shown that ORFs can also arise *de novo* from non-coding regions^24^ and that random ORFs can rapidly evolve into functional proteins^25^, offering an avenue for gene birth in a different codon size should the appropriate qtRNAs be available. However, even in an organism that had accumulated several functional qtRNAs, meaningful *de novo* quadruplet ORFs are less likely to arise by chance than equivalent triplet ORFs due to the expanded number of possible codons (256 vs. 64) and the reduced initial number of sense codons (8 vs. 61). Together, these factors contribute to the absence of naturally occurring quadruplet codon codes, despite the frequency with which individual quadruplet translation components can arise through evolution.

Although these factors explain the absence of naturally occurring all-quadruplet genetic codes, our results offer a clear blueprint for deliberately engineering an expanded all-quadruplet genetic code. With 256 total codons, this genetic code would create hundreds of free codons that could be assigned to noncanonical amino acids (ncAAs), valuable chemical additions to the genetic code that enable improved protein therapeutics^7^. Our data has revealed, for the first time, exactly which triplet codons can be reassigned to quadruplet codons without compromising amino acid selectivity: an initial all-quadruplet code will be composed of 9 selectively charged qtRNAs from this study (qtRNA^Gln^_CAGG_, qtRNA^Pro^_CCGG_, qtRNA^Phe^_CGGC_, qtRNA^Glu^_CGGT_, qtRNA^Gly^_GGGG_, qtRNA^Ser^_TCGG_, qtRNA^Arg^_AGGG_, and either qtRNA^Tyr^_TAGA_ or qtRNA^His^_TAGA_). Further, the flexibility of AARSs suggests that anticodon reengineering may be a viable strategy for reassignment of the additional 11 canonical amino acids to quadruplet codons. We and others^26^ have observed that ArgRS is a major source of promiscuous charging; refactoring or replacing this ArgRS may be broadly valuable for enabling reassignment of arginine-like codons; i.e. enabling selective charging of qtRNA^Ala^_AGGG_. Additionally, strategies such as CCA tail engineering^27,28^ will be required to mitigate toxicity and competition with triplet tRNAs. Together, our deliberate exploration of the evolution of quadruplet translation has provided components that will launch synthetic efforts to assemble a 256-amino acid genetic code.

## Acknowledgements

We thank Eric Spooner and the Whitehead Proteomics Core Facility. We thank the laboratory of Kristala Prather for equipment use and assistance. We thank Nili Ostrov, Ben Thuroni, Stephen Von Stetina, and Sergey Melnikov for their helpful comments on the manuscript. We thank BiologicsCorp for protein purification.

## Funding

This work was supported by the MIT Media Lab, an Alfred P. Sloan Research Fellowship (to K.M.E.), gifts from the Open Philanthropy Project and the Reid Hoffman Foundation (to K.M.E.), and the National Institute of Digestive and Kidney Diseases (R00 DK102669-01 to K.M.E.), and by the National Institute of General Medical Sciences (R35GM122560 to D.S.). E.A.D. was supported by the National Institute for Allergy and Infectious Diseases (F31 AI145181-01).

## Author contributions

E.A.D conceived the study. E.A.D designed the experiments with advice from K.M.E. E.A.D performed the experiments. K.M.E supervised the research. E.A.D wrote the manuscript with input from all authors.

## Competing interests

E.A.D and K.M.E have filed a patent application with the US Patent and Trademark Office on this work.

## Data and materials availability

Key plasmids from this study have been deposited on Addgene.

## Methods

### General methods

#### Antibiotics

Antibiotics (Gold Biotechnology) were used at the following working concentrations: carbenicillin, 50 μg/mL; spectinomycin, 100 μg/mL; chloramphenicol, 40 μg/mL; kanamycin, 30 μg/mL; tetracycline, 10 μg/mL; streptomycin, 50 μg/mL.

#### Media

David Rich Medium (DRM)^29^ (US biological, #CS050H-001 and #CS050H-003) is used for luminescence assays due to its low fluorescence and luminescence background. 2XYT media (US biological, T9200), a media optimized for phage growth, is used for all other purposes, including phage-based selection assays and general cloning. Agar (US biological, A0930) is used for cloning and plaque assays.

#### Plasmid cloning

tRNA genes were amplified directly from *E. coli* genomic DNA; see Table S1 for tRNA sequences and links to Benchling plasmid maps. Plasmids were cloned using either Mach1, Turbo, DH5a, or 10beta cells. Unless otherwise noted, plasmids and phage were cloned by USER assembly^30^ using the Phusion U Hot Start DNA polymerase (Thermofisher, F556L) and USER enzyme (New England Biolabs, M5505L).

#### Preparation of chemically competent cells

Strain S2060 (Addgene #105064), a K12 derivative optimized for directed evolution^31^ was used in all luciferase, phage propagation, and plaque assays. To prepare competent cells, an overnight culture was diluted 1,000-fold into 50 mL of 2XYT media supplemented with maintenance antibiotics and grown at 37 °C with shaking at 230 rpm to OD_600_ ~0.4-0.6. Cells were pelleted by centrifugation at 6,000× *g* for 10 min at 4 °C. The cell pellet was then resuspended by gentle stirring in 5 mL of TSS (LB media supplemented with 5% v/v DMSO, 10% w/v PEG 3350, and 20 mM MgCl_2_). The cell suspension was stirred to mix completely, aliquoted and flash-frozen in liquid nitrogen, and stored at −80 °C until use.

#### Transformation of chemically competent cells

To transform cells, 100 μL of competent cells were thawed on ice. To this, plasmid (2 μL each of miniprep-quality plasmid; up to two plasmids per transformation) and 100 μL KCM solution (100 mM KCl, 30 mM CaCl_2_, and 50 mM MgCl_2_ in H_2_O) were added and stirred gently with a pipette tip. The mixture was incubated on ice for 10 min and heat shocked at 42 °C for 45 s. The mixture was chilled on ice for 4 min, then 850 μL of 2XYT media was added. Cells were allowed to recover at 37 °C with shaking at 230 rpm for 0.75 h, streaked on 2XYT media + 1.5% agar plates containing the appropriate antibiotics, and incubated at 37 °C for 16–18 h.

#### Standard phage cloning

Competent *E. coli* S2060 cells were prepared containing pJC175e (Addgene #79219), a plasmid expressing pIII under control of the phage shock promoter^32^, which enables propagation of ΔpIII M13 bacteriophage through complementation. To clone ΔpIII M13 bacteriophage, PCR fragments were assembled using USER. The annealed fragments were transformed into competent S2060-pJC175e competent cells. Transformants were recovered in 2XYT media overnight, shaking at 230 rpm at 37 °C. The phage supernatant from the resulting culture was filtered through a 0.22 μm membrane (Thomas Scientific, 1166U41), and plaqued to isolate clonal phage (see below). Clonal plaques were picked into 2mL of 2XYT media and expanded overnight, shaking at 230 rpm at 37 °C, filtered, and Sanger sequenced.

#### tRNA diagrams

R2R was used to generate tRNA diagrams^33^. R2R is free software available from http://www.bioinf.uni-leipzig.de/~zasha/R2R/.

### Phage plaque assays

#### Manual protocol

S2060 cells were transformed with the Accessory Plasmid (AP) of interest. Overnight cultures of single colonies grown in 2XYT media supplemented with maintenance antibiotics were diluted 1,000-fold into fresh 2XYT media with maintenance antibiotics and grown at 37 °C with shaking at 230 rpm to OD_600_ ~0.6-0.8 before use. Bacteriophage were serially diluted 100-fold (4 dilutions total) in H_2_O. 100 μL of cells were added to 100 μL of each phage dilution, and to this 0.85 mL of liquid (70 °C) top agar (2XYT media + 0.6% agar) supplemented with 2% Bluo-Gal (GoldBio, CAS #97753-82-7) was added and mixed by pipetting up and down once. This mixture was then immediately pipetted onto one quadrant of a quartered petri dish (VWR, 25384308).already containing 2 mL of solidified bottom agar (2XYT media + 1.5% agar, no antibiotics). After solidification of the top agar, plates were incubated at 37 °C for 16-18 h.

#### Robotics-accelerated protocol

Plaque assays were automated as previously described^34^. Briefly, the same procedure was followed as above, except that plating of the plaque assays was done by a liquid handling robot (Hamilton Robotics) by plating 20 μL of bacterial culture and 100 μL of phage dilution with 200 μL of soft agar onto a well of a 24-well plate already containing 235 μL of hard agar per well. To prevent premature cooling of soft agar, the soft agar was placed on the deck in a 70 °C heat block.

### Luciferase readthrough assay

Luciferase readthrough assays were performed as previously described^36^. Briefly, S2060 bacteria were transformed with a luciferase reporter containing a quadruplet codon at residue 357 (for example, https://benchling.com/s/seq-7CsWcP8Ez4JNjNM9W23N, Addgene #134787) and an inducible qtRNA expression plasmid (for example, https://benchling.com/s/seq-F5NZDNmxOhUoWp5DX41h, Addgene #134800). Bacteria were grown overnight, then diluted 500-fold the next day into DRM media with qtRNA expression induced (1mM IPTG, GoldBio, I2481C5), or suppressed (0mM IPTG). Absorbance and luminescence measurements are taken kinetically in a ClarioSTAR plate reader (BMG Labtech) over the course of 8 h. To account for differential growth rate, all luminescence values are considered at OD_600_ = 0.5. The % of Triplet translation efficiency is calculated using the formula (QuadLux _qtRNA induced_ – QuadLux _qtRNA uninduced_)/ (TriLux – QuadLux _qtRNA uninduced_), where:

– TriLux is the luminescence of the positive control, a luciferase encoded entirely with triplet codons
– QuadLux _qtRNA induced_ is the luminescence produced by the quadruplet codon-bearing reporter upon qtRNA expression (1 mM IPTG)
– QuadLux _qtRNA uninduced_ is the luminescence produced by the quadruplet codon-bearing reporter in the absence of qtRNA expression (0 mM IPTG)

### Growth defect

Our measurement of growth defect involves analysis of the absorbance measured during the 8h growth curves taken in the luciferase readthrough assay. We identify the time, t_measure_ at which QuadLux _qtRNA induced_ reaches OD_600_ = 0.5. Growth defect = 0.5/ (OD of QuadLux _qtRNA suppressed_ at t_measure_).

### Phage library primer design

We do not recommend USER cloning for library creation inside of high-secondary structure tRNAs; instead, we used degenerate primers and blunt end ligation. Primers were designed for use with around-the-world PCR, creating a one-piece blunt-end ligation. In order to reduce nucleotide bias during blunt end ligation assembly, the last degenerate base was designed to be at least one base away from the end of the primer. In all cases, phage bearing the wildtype triplet tRNA scaffold are used as a template for the PCR in order to eliminate background bias in the libraries.

#### Degenerate NNNN anticodon libraries

Degenerate nucleotides are included at bases 34, 35, 35.5 and 36 of the qtRNA. See an example of this primer design: https://benchling.com/s/seq-5k2FZdPbWpiB8tOh8WCK

#### Primer design library diversified at positions 32, 37 and 38 of qtRNA

Degenerate nucleotides are included at bases 32, 37 and 38 of the qtRNA. See an example of this primer design: https://benchling.com/s/seq-oMMGFRl2vITYpNX2HWkz

### Phage library cloning

Primers for around-the-world PCR and blunt end ligation were designed as described above. For each library, 200 μL of PCR product was produced using Phusion Hotstart Flex polymerase (New England Biolabs, M0535S). The entirety of this PCR product was run on a gel, extracted, and purified using spin column purification (Qiagen, 28106). Background plasmid was digested using Dpn1 (New England Biolabs, R0176L), and the remaining PCR product was purified again using spin columns, and ligated. The ligation product was transformed into competent E. coli S2060 cells containing pJC175e (as described). Transformants were recovered in 2XYT media overnight, shaking at 230 rpm at 37 °C. The phage supernatant from the resulting culture was filtered through a 0.22 μm membrane (Thomas Scientific, 1166U41), and plaqued (as described).

### pIII-based selection of NNNN anticodon libraries

Competent S2060 cells were transformed with a plasmid expressing the appropriate quadruplet codon replacing permissive residue pIII-29^29^. A picked colony was grown overnight in 2XYT media with maintenance antibiotics. Following overnight growth, cultures were diluted 1:1000 in 2XYT media without antibiotics and 500 μL of diluted culture was aliquoted into a 2 mL deep 96-well plate. Wells were inoculated with phage encoding the desired qtRNA library to a final concentration of 10^5^ pfu/mL. The plate was grown overnight, shaking at 230 rpm at 37 °C, and then the phage supernatant was filtered and plaqued in activity-independent host S2060 cells bearing pJC175e. Selections were ranked from lowest final phage titer to highest final phage titer. Two plaques per selection were picked for Sanger sequencing in all selections that enrich over 10-fold, or additional selections as desired.

### pIII-based selection and NGS of libraries diversified at positions 32, 37 and 38

Competent S2060 cells were transformed with a plasmid expressing the appropriate quadruplet codon replacing permissive residue pIII-29^29^. A picked colony was grown overnight in 2XYT media with maintenance antibiotics. Following overnight growth, cultures were diluted 1:1000 in 2XYT media without antibiotics and 500 μL of diluted culture was aliquoted into a 2 mL deep 96-well plate. Wells were inoculated with phage encoding the desired qtRNA library to a final concentration of 10^5^ pfu/mL. The plate was grown overnight, shaking at 230 rpm at 37 °C, and then the phage supernatant was filtered and plaqued in activity-independent host S2060 cells bearing pJC175e. Libraries were amplified using PCR and sequenced on a MiSeq to >10x library size.

### Quantification of qtRNA charging using mass spectrometry

Each qtRNA (for example, https://benchling.com/s/seq-BpqbupgoHvZu0gFmNUBm) was co-expressed with C-terminal 6xHis-tagged *sfGFP* (for example, https://benchling.com/s/seq-bI1bixktGKegGwboMYIP) with the appropriate quadruplet codon replacing permissive residue 151 in S2060 cells. GFP was purified from these cultures at either 5mL or 1L scale (Table S2).

#### Purification at 4mL scale

Co-transform bacteria with sfGFP-151-quad and qtRNA expression plasmids. Inoculate a single colony of the expression strain in DRM media (with 100 ug/ml spectinomycin, 30 ug/ml kanamycin, and 25 mM glucose) as the seed culture; grow overnight at 37°C, 200 rpm. Dilute 1:1000 and into DRM media that induces qtRNA expression (with 100 ug/ml spectinomycin, 30 ug/ml kanamycin, and 1 mM IPTG) and grow for 29 hours, 200rpm, at 37C. Cultures were spun down at 5000g for 10 minutes and the pellet frozen at −80C. Thawed pellets were lysed using B-PER Complete Reagent (Thermofisher 89821) and His-tagged protein was purified from cell lysate using a Ni-NTA spin column (Qiagen, 31014). The resulting product was run on a 12% Bis-Tris PAGE gel (Thermofisher, NP0342PK2), and the appropriate band was extracted for mass spec analysis.

#### Purification at 1L scale

Proteins were purified by BiologicsCorp at 1L scale. Inoculate a single colony of the expression strain in LB media (with 100 ug/ml spectinomycin, 30 ug/ml kanamycin, and 25 mM glucose) as the seed culture; at 37°C, 200 rpm, overnight. Perform 1% inoculation into 500 ml TB media (with 100 ug/ml spectinomycin, 30ug/ml kanamycin, and 25 mM glucose); at 37°C, 200 rpm, till OD_600_ = 0.6-0.8. Add IPTG to a final concentration of 1 mM. Continue culturing for 4 hrs. Collect cells by centrifugation (4°C, 8,000 rpm, 15 min). Wash cells once by resuspension with PBS buffer and again collect cells by centrifugation. Resuspend per gram of wet cell pellet with 20 ml Lysis Buffer T300-20L (50 mM Tris-HCl pH 8.0, 300 mM NaCl, and 20 mM imidazol, plus 1 mM DTT, 1% Triton X-114, 1 ug/ml Pepstatin A, and 1 ug/ml Leupeptin). Sonicate the cell suspension on ice for 15-25 min (500 W; 3 sec burst and 6 sec pause). Centrifuge the cell lysate; at 4C, 12500 rpm, for 15 minutes. Filter the supernatant through a 0.45 um pore membrane. Purify His-tagged protein using a Ni-IDA column. Analyze fractions by SDS-PAGE. For samples with satisfactory purity, pool fractions and dialyzed against PBS buffer (pH 7.4); at 4C; 2 hr per cycle, 3-4 cycles. Filter the protein solution through a 0.22 um pore membrane. Concentrate if necessary.

#### Mass spectrometry analysis

Samples were trypsin digested and measured using HPLC-MS/MS. To analyze the results, the resulting fragmentation spectra were correlated against a custom database containing the predicted fragmentation pattern for the fragment straddling residues 151 if it contained each of the 20 canonical amino acids. Abundance of each species was quantified by calculating the area under the curve of the ion chromatogram for each peptide precursor. The limit of detection was 10^4^ [AU], the lower limit for area under the curve for a peptide on this instrument.

---

**Figure S1.**
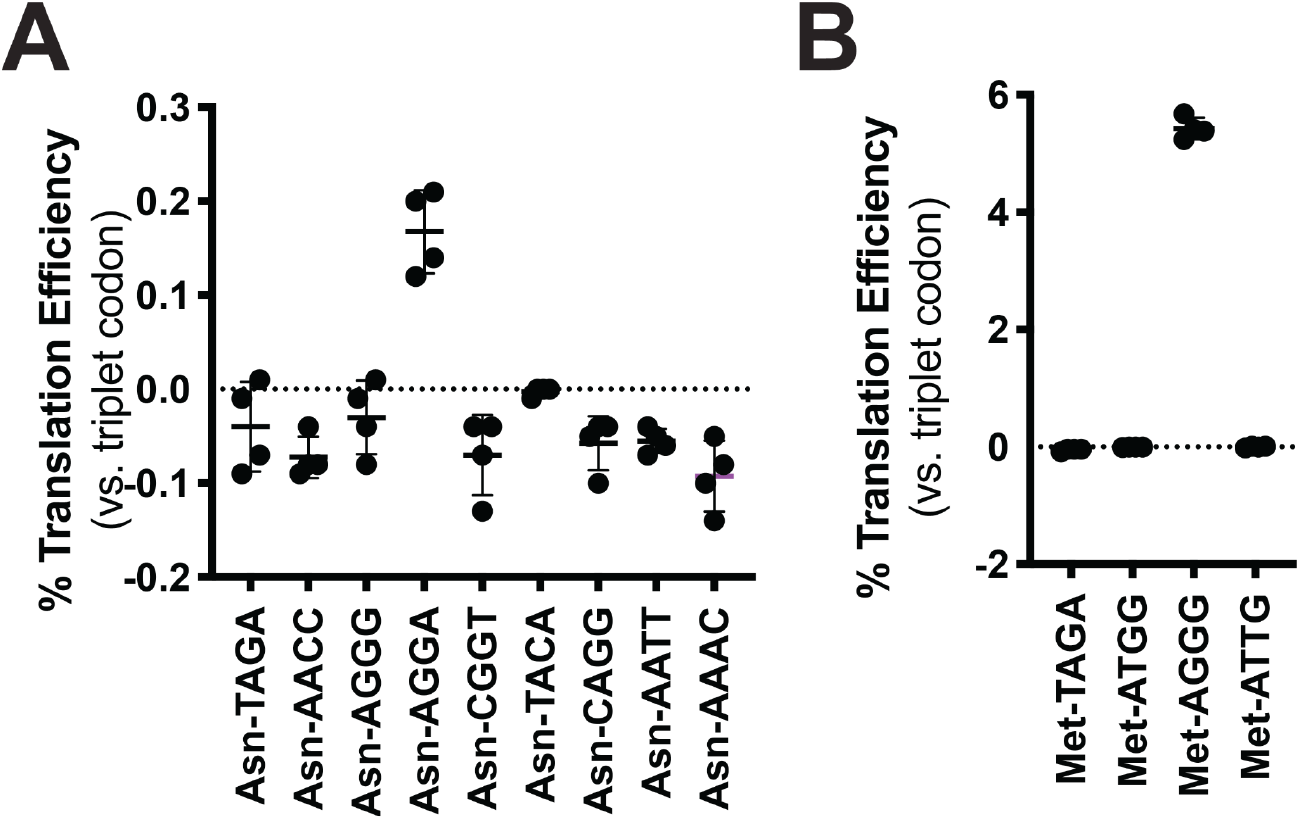
Codon reassignment for asparagine and methionine. Quadruplet codon experimentation for asparagine and methionine. Measurements are taken kinetically and normalized to culture density. Efficiency is reported relative to luminescence produced by a WT, all triplet luciferase transcript. **A)** Characterization of nine unique quadruplet codons within the tRNA^Asn^ scaffold. **B)** Characterization of four quadruplet codons within the tRNA^Met^ scaffold.

**Table S1.**
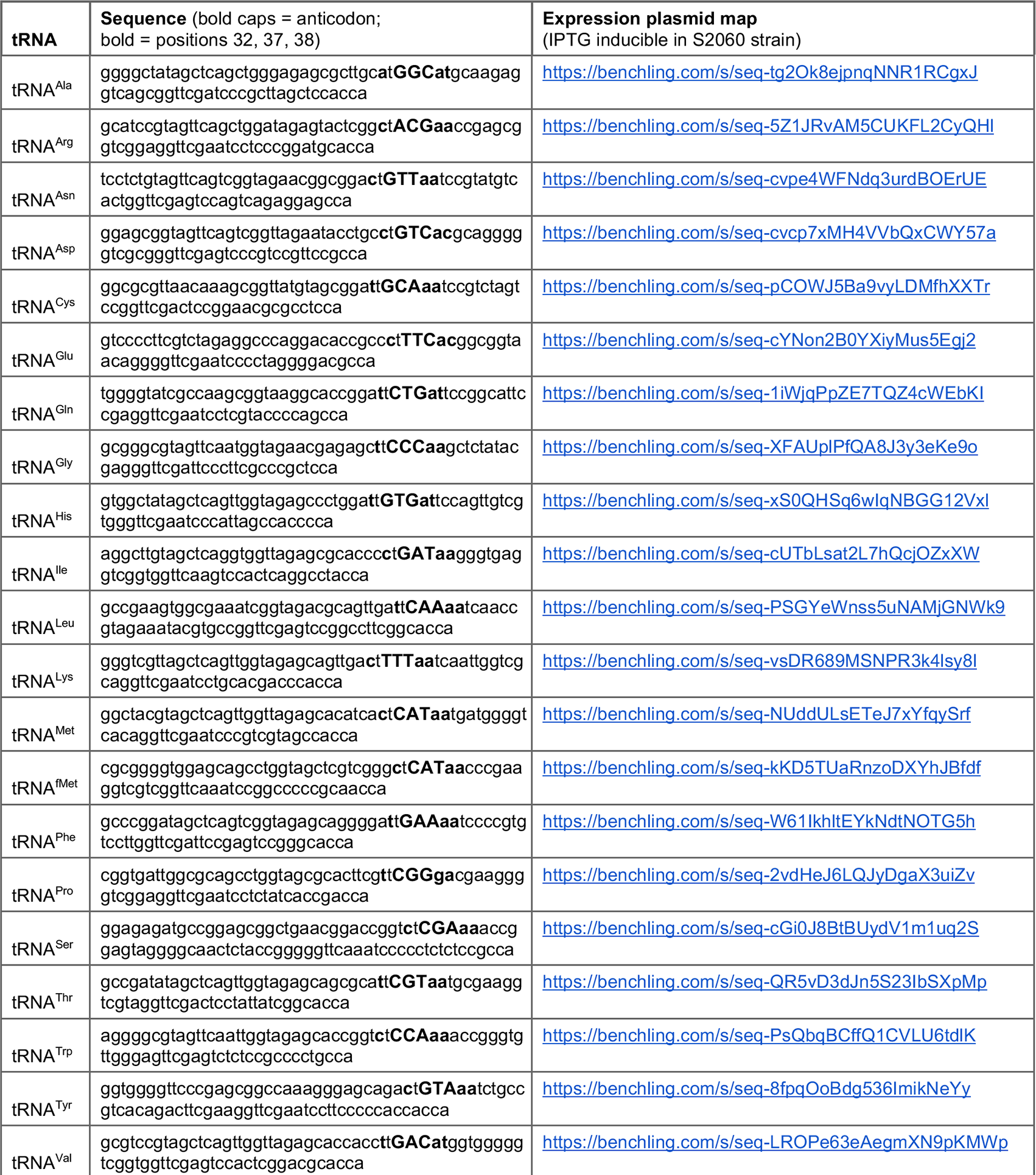
Table of tRNA scaffolds. Canonical triplet tRNAs were cloned from *E. coli* K12 genomic DNA. Body text Figures 1 and 2 concern mutations to the anticodon (bolded caps); body text Figure 3 concerns mutation to positions 32, 37 and 38 (bolded lower case). We use qtRNA^three letter scaffold^_four letter DNA codon_ nomenclature to refer to qtRNAs; e.g. to create qtRNA^Ser^_TAGA_, the CGA anticodon in tRNA^Ser^ scaffold would be replaced by TCTA.

**Table S2.**
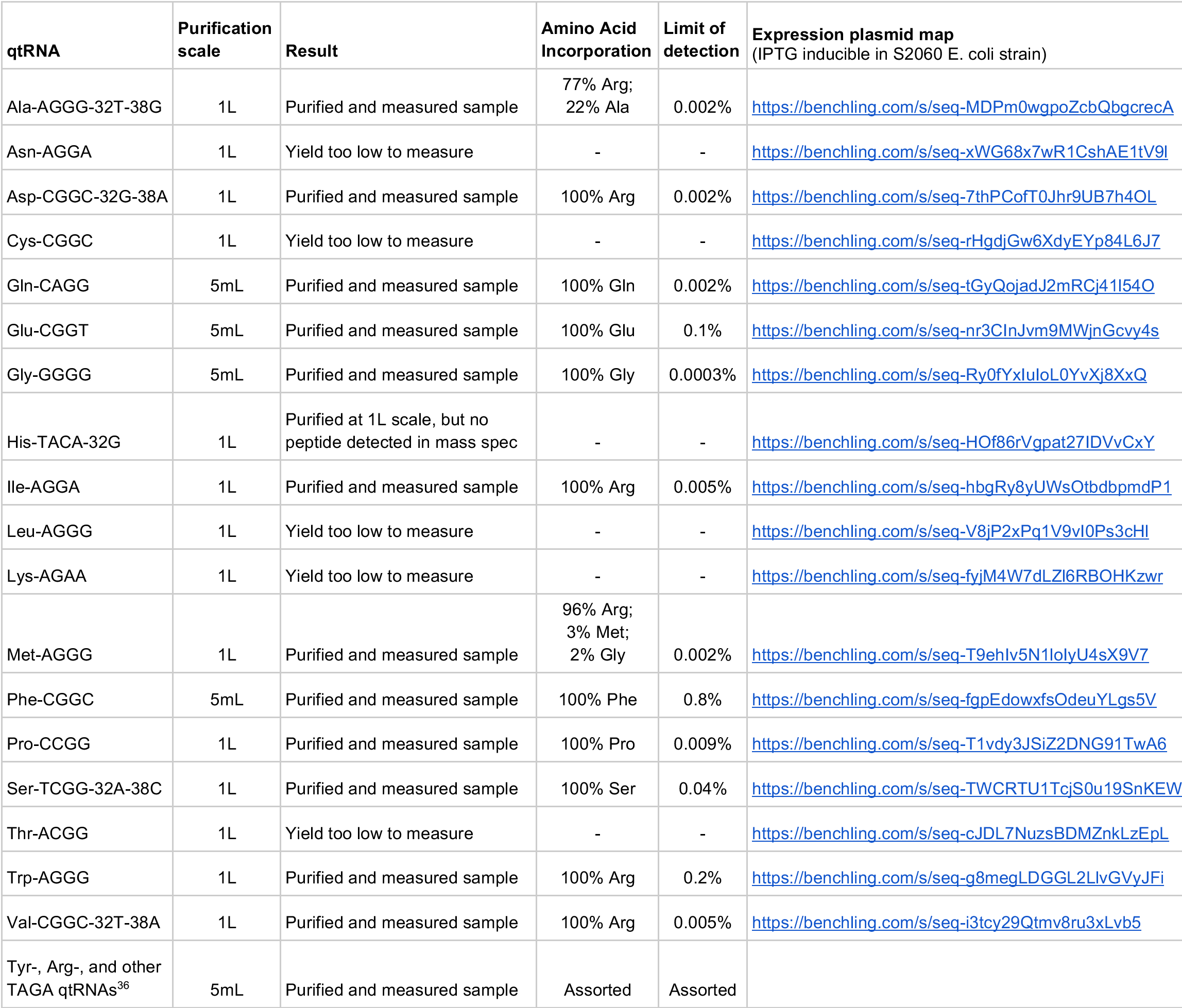
purification scale of qtRNA-translated proteins. For sfGFP purification, qtRNA expression plasmids (listed above) were co-expressed with C-terminal 6xHis-tagged *sfGFP* with the appropriate quadruplet codon replacing permissive residue 151. For example, sfGFP-151-GGGG, https://benchling.com/s/seq-bI1bixktGKegGwboMYIP

## Notes

### Competing Interest Statement

The authors have declared no competing interest.

